# New inhibitors of the *Pseudomonas aeruginosa* enzyme, PqsE, and methods assessing their potential to induce a conformational change via active site binding

**DOI:** 10.1101/2025.10.07.680923

**Authors:** Samantha B. Orr, Hannah A. Jones, Margaret G. O’Hara, Kaitlyn R. Smith, Isabelle R. Taylor

## Abstract

*Pseudomonas aeruginosa* is an opportunistic pathogen known for its ability to produce virulence factors and form biofilms. These, among many other traits, enable *P. aeruginosa* to cause infections, evade host immune responses, and resist treatment with antimicrobial agents. Both the ability to form biofilms and produce virulence factors are regulated via the bacterial cell-cell communication process called quorum sensing. A key molecular event that enables quorum sensing in *P. aeruginosa* is the physical interaction between an enzyme, PqsE and a master quorum-sensing receptor/transcription factor RhlR, which regulates the expression of a wide variety of virulence-associated genes. Previous work identified active site mutations in PqsE that induce a conformational change weakening the interaction with RhlR. These active site mutations weakened the PqsE-RhlR interaction to the extent that several virulence genes were not activated, and the mutant strains of *P. aeruginosa* failed to colonize the lungs of a mouse. In this study, we designed a series of molecules to probe binding in the active site of PqsE as a strategy for inhibiting the PqsE-RhlR interaction and thus, the activation of RhlR-controlled genes in *P. aeruginosa*. HJ1 and HJ5 are new molecules that both bind in the active site of PqsE, and while HJ5 appears to bind in an alternate mode compared to HJ1, it does not induce a conformational change to weaken the PqsE-RhlR interaction. Here, we introduce multiple experimental approaches to assess the way in which these new molecules engage in the PqsE active site. HJ5 can serve as a promising starting point for the development of molecules that target the PqsE active site and allosterically inhibit the interaction with RhlR, thus decreasing virulence in *P. aeruginosa*.

## Introduction

*Pseudomonas aeruginosa* is an opportunistic bacterial pathogen primarily responsible for causing infections in the immunocompromised population, especially those with cystic fibrosis (1–3). *P. aeruginosa* is highly resistant to a wide range of antibiotics and poses a serious threat to public health (4–6). Therefore, *P. aeruginosa* has been classified by both the World Health Organization and Centers for Disease Control and Prevention as a high priority for the development of antibiotics with novel mechanisms of action (7, 8). Among the many pathogenic behaviors that make *P. aeruginosa* such a threat are the ability to form biofilms and produce virulence factors (9, 10). Expression of the genes involved in these behaviors is controlled by a cell-to-cell communication process called quorum sensing (11, 12).

Quorum sensing (QS) is the process through which bacterial cells monitor both population density and species composition in the surrounding environment (13, 14). This is achieved by the synthesis, release, and detection of small molecule signals called autoinducers (15). Each autoinducer is produced by a dedicated enzymatic synthase, or a biosynthetic pathway, and binds a cognate receptor/transcription factor in order to regulate gene expression in response to cell density. One branch of the *P. aeruginosa* QS system consists of a synthase, RhlI, that produces the autoinducer *N*-butyryl-homoserine lactone (C4-HSL) which binds to the receptor, RhlR. Binding of C4-HSL activates RhlR as a transcription factor, regulating a wide range of virulence-related genes (16, 17). Notably, the ability of RhlR to act as a transcription factor is additionally dependent on a protein-protein interaction with PqsE, a redundant metallo-β-hydrolase from the *Pseudomonas* Quinolone Signaling (PQS) system (18–20).

The ability of PqsE to form a complex with RhlR is independent of PqsE catalytic function, however particular mutations in the active site have been shown to induce a conformational change that severely weakens the affinity of PqsE for RhlR (18, 21). These active site mutations that weaken the PqsE-RhlR interaction disrupt the production of virulence factors, such as the blue-green toxin pyocyanin, and decrease colonization of *P. aeruginosa* in the lungs of a mouse (21, 22). These findings strongly suggest that a small molecule disruptor of PqsE-RhlR complex formation could be an effective *P. aeruginosa*-targeted antibiotic. Our goal is to develop new small molecules that bind in the active site of PqsE, and induce a conformational change similar to that observed in the PqsE(E182W) variant. The E182W substitution was shown to induce a loop rearrangement that repositions another residue, E280 into the active site (21). The exact mechanism by which the loop rearrangement weakens the affinity of PqsE(E182W) for RhlR is not well understood, but it is suspected to inhibit PqsE dimerization, which is necessary for proper complex formation with a functional RhlR dimer (23–25). Our goal is to find a molecule that binds in the active site of PqsE and forces a conformational change resembling that of the PqsE(E182W) variant.

Starting with a small molecule that exhibits potent inhibition of PqsE enzyme activity and for which a crystal structure had been previously solved (21), we designed and synthesized a series of active site-targeted PqsE inhibitors. One derivative, HJ5, was designed to interact with the most buried portion of the PqsE active site, where the E182 sidechain resides. This derivative does retain the ability to bind in the active site and inhibit catalytic function of PqsE, but appears to have a differential inhibition profile compared to the parent derivative, HJ1. While HJ5 does not cause a conformational change in PqsE, or inhibit the PqsE-RhlR protein-protein interaction, this small molecule series is a new starting point for the development of PqsE inhibitors with the potential to allosterically disrupt PqsE-RhlR complex formation. We also present new assessments for determining the effect of small molecules on PqsE conformational dynamics and PqsE-RhlR-driven virulence in *P. aeruginosa*.

## Results

### A new scaffold, HJ1, has similar potency to BB584

BB584 was originally synthesized along with a series of other derivatives designed to assess the importance of particular moieties for strong binding affinity in the PqsE active site (21). BB584 was determined to be the tightest binding derivative of the series, and a crystal structure of the PqsE-BB584 complex was solved. From the crystal structure, it was confirmed that the indazole ring is oriented at the most buried portion of the PqsE active site, and therefore, presents the most ideal part of the molecule to derivatize in order to potentially induce a conformational change similar to the one observed in the PqsE(E182W) variant (**Figure 1a**). The tert-butyl group of BB584 is positioned at the most solvent-exposed portion of the active site and seemed the least important for binding. We noted that the entire tert-butyl-substituted urea resembled a tert-butyloxycarbonyl (boc) protecting group, and that based on the crystal structure, replacing this group with a boc group would likely preserve all important binding interactions of a small molecule in the active site. This change would also greatly simplify the synthesis of such a derivative, and therefore, HJ1 was designed and synthesized in one amide coupling step with commercial starting materials (**Figure 1a**). When tested for its ability to inhibit enzyme function of PqsE, HJ1 was found to have similar, although slightly decreased potency compared to BB584 (IC_50_ of HJ1 = 264 nM, IC_50_ of BB584 = 108 nM) (**Figure 1b**). The similar potency, and therefore binding affinity, of HJ1 for the PqsE active site compared to BB584 encouraged us to use this synthetically simplified scaffold as a starting point for the synthesis of new active site-targeted derivatives.

**Figure 1:**
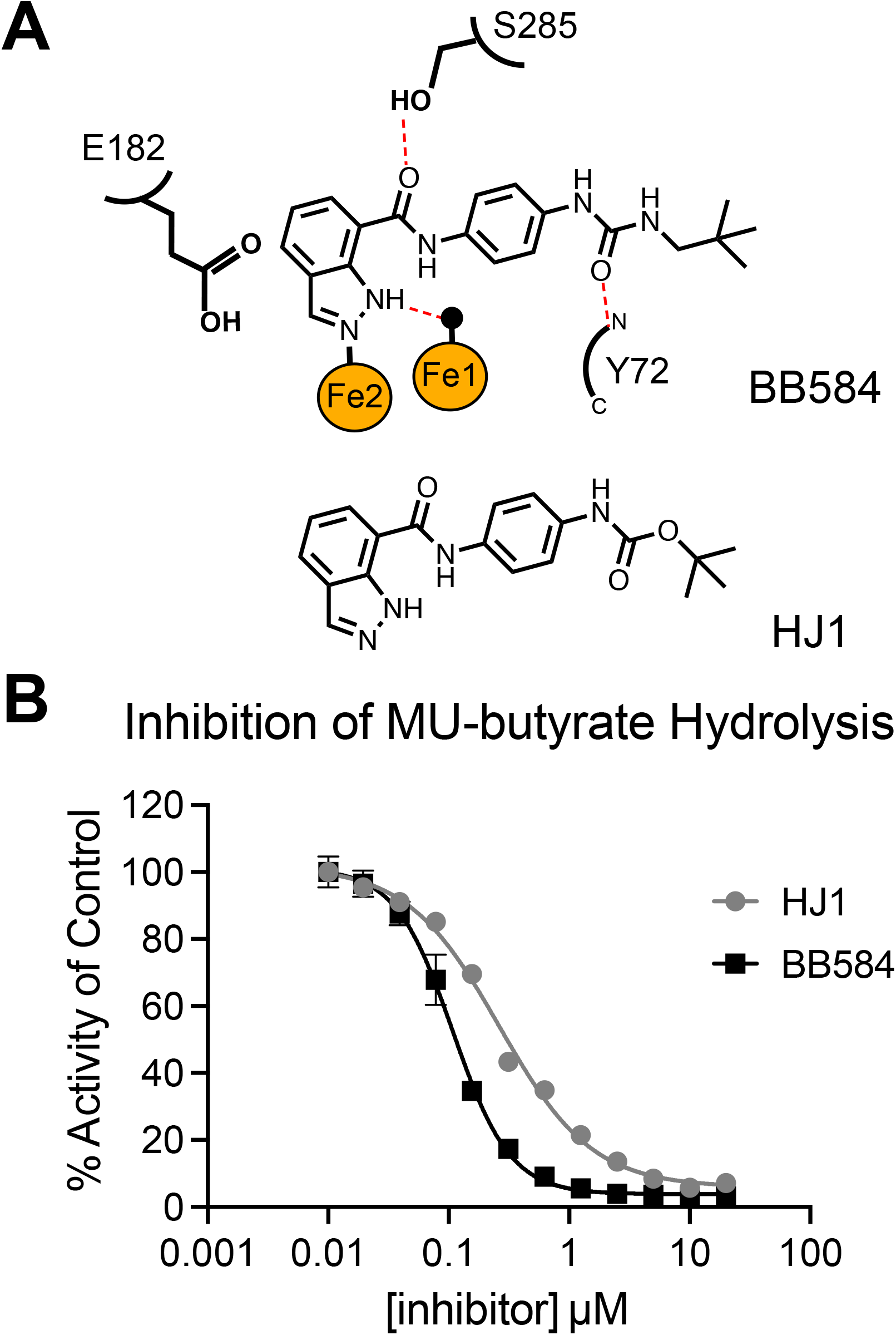
Design of HJ1 and potency compared to BB584. A) Outlined binding pose of BB584 in the active site of PqsE with key active site residues highlighted (derived from PDB: 7TZA). Hydrogen bonds are shown as red dashed lines, an active site water is shown as a black sphere, and the two iron atoms are shown as orange spheres. The structure of HJ1 is shown below BB584 for comparison. B) Inhibition of PqsE-catalyzed MU-butyrate hydrolysis by BB584 and HJ1. Background fluorescence was measured for the compound dilution series plus MU-butyrate in the absence of PqsE and was subtracted from the values plotted prior to normalization. The data were normalized so that DMSO-treated PqsE is 100 % activity. Data shown are the average of technical triplicates and error bars represent standard deviation.

### Synthetic derivatization of HJ1 extends small molecule interaction into the PqsE active site

Two positions on the indazole ring of HJ1 were determined as the starting points for synthetic derivatization due to their predicted proximity to the E182 residue. The first of these positions (highlighted in blue in **Figure 2**) was methylated to give the product, HJ2. When tested in an esterase assay, this derivative was found to have virtually no effect on enzymatic activity of PqsE, and therefore, likely lost all binding affinity with the introduction of the methyl group (**Figure 2**). Derivatives featuring a chloro or bromo substituent at this position were also synthesized, and additionally found to completely lack inhibitory activity (HJ3 and HJ4, **Figure S1**).

**Figure 2:**
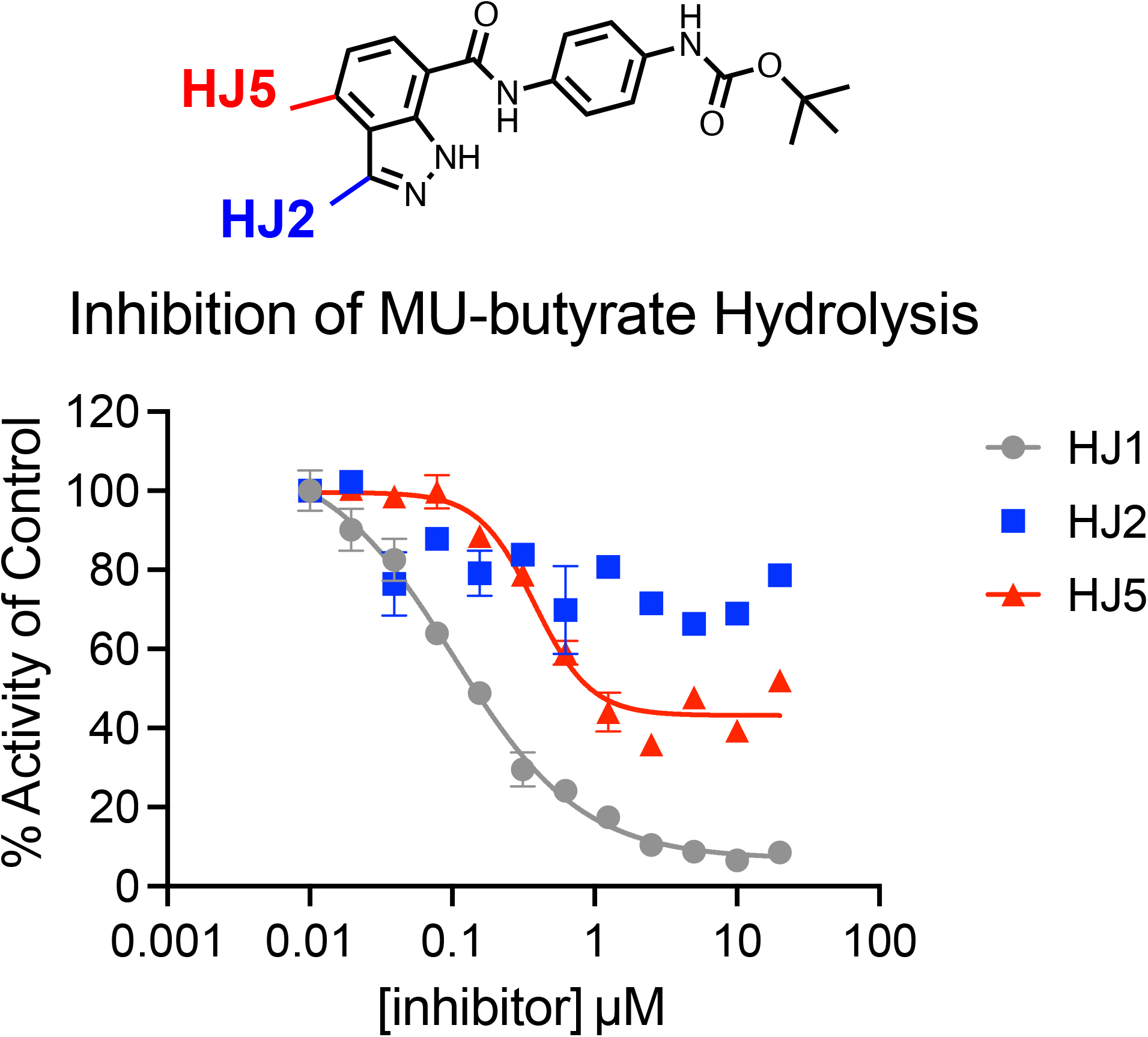
Methylated derivatives of HJ1 and their inhibitory activity. Two positions on the indazole ring of HJ1 were methylated to give the synthetic derivatives HJ2 (blue) and HJ5 (red). Inhibition of PqsE-catalyzed MU-butyrate hydrolysis is shown. Background fluorescence was measured for the compound dilution series plus MU-butyrate in the absence of PqsE and was subtracted from the values plotted prior to normalization. The data were normalized so that DMSO-treated PqsE is 100 % activity. Data shown are the average of technical triplicates and error bars represent standard deviation.

We then switched focus to derivatization of the other highlighted position on the indazole ring. This derivatization required an alteration to the synthetic route, as starting carboxylic acids with substitutions at this position were not commercially available. In order to prepare the 4-substituted indazole carboxylic acid, methyl 2-amino-3,4-dimethylbenzoate was first reacted with isopentyl nitrite to form the methylated indazole ring. The resulting ester was then demethylated to give the starting carboxylic acid and an amide coupling reaction was carried out to give HJ5. Once HJ5 was isolated and purified, it was tested in an esterase inhibition assay and found to have similar potency to HJ1 (IC_50_ of HJ5 = 379 nM). Curiously, while HJ1 exhibits a full inhibition curve that brings enzymatic activity of PqsE down to baseline levels (i.e. 0% activity compared to the DMSO-treated control protein), HJ5 only inhibited esterase activity to a baseline of approximately 40-50% of the hydrolytic capacity of the uninhibited enzyme. This result seemed to suggest that the mode of inhibition or binding of HJ5 in the active site may be different from that of HJ1. We were encouraged by the finding that substitution of this position on the indazole ring of the HJ1 scaffold would still lead to binding, and explored further whether binding of HJ5 in the PqsE active site could induce a conformational change in PqsE.

### Insights to the possible binding modes of HJ1 and HJ5

The inhibitory activity of HJ5 was encouraging, although due to the incomplete inhibition profile, it was necessary to further characterize binding using an alternate method. To confirm binding of HJ5 to PqsE, we used a differential scanning fluorimetry assay to measure the shift in melting temperature (T_m_) induced by small molecule binding. This technique has been used previously to screen for small molecule PqsE inhibitors (26) and to characterize their binding to various purified PqsE variants *in vitro* (18). Purified PqsE(WT) was treated with DMSO, HJ1, HJ2, or HJ5 and the resulting T_m_ was measured. While treatment with HJ2 led to only a small shift in T_m_ (ΔT_m_ = 0.7 ± 0.2 °C), consistent with a general lack of inhibitory activity, binding of HJ1 to PqsE(WT) led to a shift of 3.6 ± 0.4 °C and HJ5 binding led to a shift of 1.8 ± 0.6 °C (**Figure 3**). These shifts in T_m_ were determined to be significant compared to the ΔT_m_ observed for HJ2-treated PqsE(WT). Based on the crystal structure of BB584 bound to PqsE, we anticipated that binding of HJ1 and HJ5 would depend largely on the ability to form a hydrogen bond with the S285 sidechain. We therefore were interested in testing whether substitution of the serine with either an alanine or a space-filling tryptophan would affect binding of these molecules. In both the case of the PqsE(S285A) and PqsE(S285W) variants, neither HJ1 nor HJ5 induced significant shifts in T_m_ compared to HJ2 (**Figure 3**). From these findings, we suspect that binding affinity for the HJ series of molecules largely stems from hydrogen bonding with the active site S285 sidechain. HJ1 and its methylated derivatives were also tested for binding to other previously characterized variants, although the results were variable and no significant differences in ΔT_m_ were observed compared to the non-binding HJ2. The trends observed in these experiments were interesting however, and will be the subject of further investigation (**Figure S2**).

**Figure 3:**
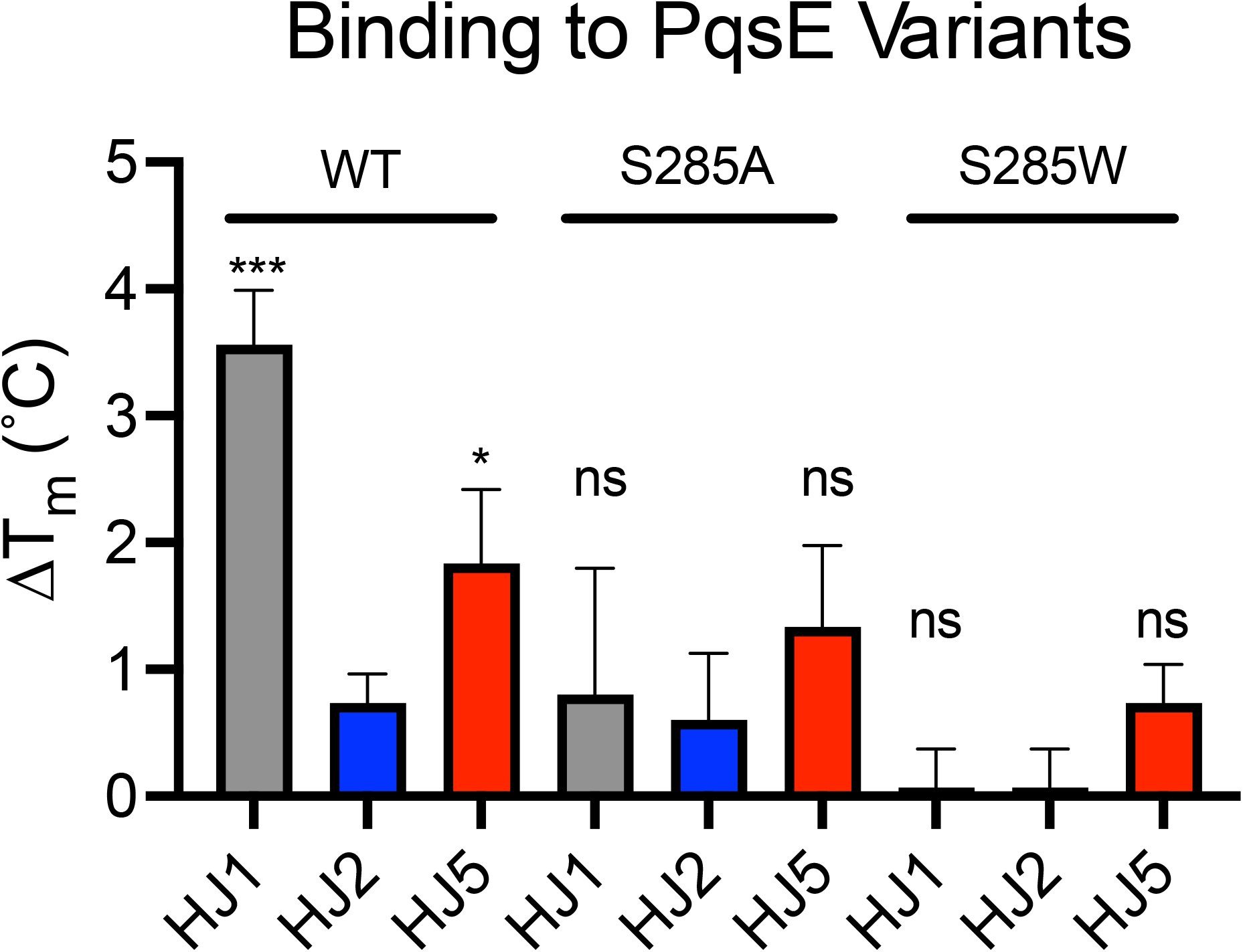
Binding of HJ derivatives to PqsE and PqsE variants. Purified PqsE proteins (2 µM) were treated with DMSO or 100 µM HJ1 (grey), HJ2 (blue), or HJ5 (red) and fluorescence of SYPRO-Orange was measured (ex: 580 nm, em: 623 nm) over 25-90 °C. The shift in T_m_ is shown compared to the DMSO-treated control. Data shown are the average of three independent experiments performed in triplicate and error bars represent standard deviation. Significance was determined via a one-way ANOVA comparing either the ΔT_m_ for HJ1 binding or HJ5 binding to the non-binding HJ2 shift (^***^ P value < 0.0005, ^*^ P value < 0.05, ns = not significant).

We were intrigued by the possibly differential effect of the S285A and S285W substitutions on binding of HJ1 and HJ5 to PqsE, so we tested the ability of these molecules to inhibit enzyme activity of the purified PqsE(S285A) and PqsE(S285W) variants in an esterase assay. Both PqsE(S285A) and PqsE(S285W) had been previously assessed for their ability to catalyze hydrolysis of the synthetic ester substrate, 4-methylumbelliferyl butyrate (MU-butyrate) *in vitro*, and had been found to possess equal or better enzymatic capacity compared to PqsE(WT) (18). While the S285W substitution partially blocks the active site, and has been shown to affect the binding affinity of small molecules and potency of active site-targeted inhibitors of PqsE, this substitution, curiously, seems to have little effect on catalytic turnover of MU-butyrate. This mutation has also been shown to have no effect on PqsE-RhlR-dependent production of the toxin, pyocyanin, in *P. aeruginosa*. When tested for dose-dependent inhibition of enzymatic activity, the potency of both HJ1 and HJ5 decreased dramatically against the PqsE(S285A) variant, with each IC_50_ increasing approximately 10-fold (**Figure 4a&b**). The maximum level of inhibition also decreased, where HJ1 went from 100% inhibition of PqsE(WT) to inhibiting only ∼70% activity of PqsE(S285A) (HJ5 went from ∼50% inhibition of PqsE(WT) to ∼10% inhibition of PqsE(S285A)). This suggests that a large degree of the affinity each of these molecules has for binding in the PqsE active site comes from the hydrogen bond formed with the serine hydroxyl group at this position. The space-filling capacity of the S285W substitution, however, completely abrogated binding of both HJ1 and HJ5 in the active site. Because the S285W substitution had such a dramatic effect on binding of these molecules in the active site of PqsE and this mutation has been previously shown to have no effect on formation of the PqsE-RhlR complex, this mutant could be used to determine whether any *in vivo* activity of HJ1 or HJ5 could be attributed to their binding in the PqsE active site.

**Figure 4:**
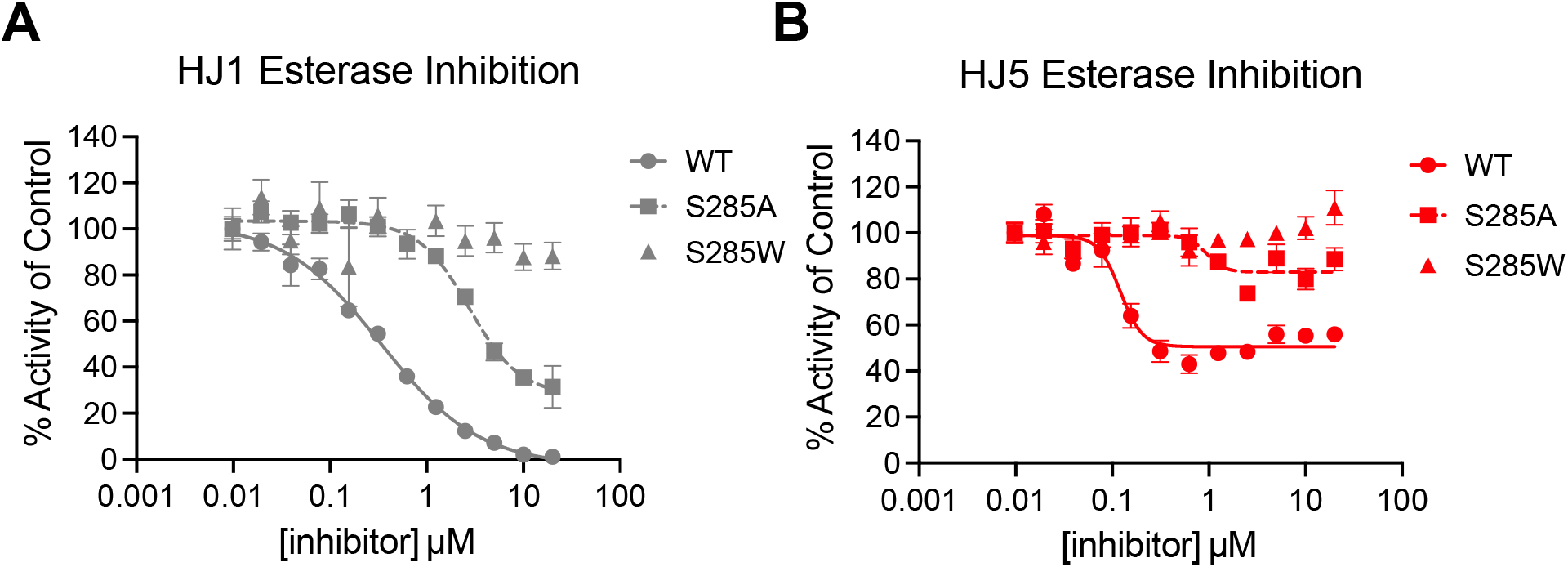
Amino acid substitutions at S285 block binding of HJ1 and HJ5 in the PqsE active site. A) HJ1 inhibition and B) HJ5 inhibition of MU-butyrate hydrolysis by PqsE(WT) (circles), PqsE(S285A) (squares), and PqsE(S285W) (triangles). Background fluorescence was measured for the compound dilution series plus MU-butyrate in the absence of PqsE and was subtracted from the values plotted prior to normalization. The data were normalized so that DMSO-treated PqsE is 100 % activity. Data shown are the average of technical triplicates and error bars represent standard deviation.

### HJ5 decreases pyocyanin production through an off-target mechanism

The observation that HJ1 and HJ5 are incapable of binding the PqsE(S285W) mutant was an exciting finding, because it presented a way for us to test whether these molecules could inhibit PqsE-dependent virulence phenotypes in *P. aeruginosa* via an on-target mechanism. In particular, pyocyanin production is dependent on the PqsE-RhlR protein-protein interaction (18, 19, 27). It has been previously established that introduction of the E182W substitution in the active site of PqsE causes a conformational change in the protein that severely weakens its affinity for RhlR, and leads to a dramatic decrease in pyocyanin production (21). While the E182W mutation has this effect, the S285W mutation has no effect on either PqsE-RhlR complex formation or pyocyanin production. Therefore, if a molecule from this series is binding in the active site of PqsE and causing a conformational change that weakens affinity for RhlR, we would expect such a molecule to inhibit pyocyanin production in a WT strain of *P. aeruginosa*, but not in a strain harboring *pqsE(S285W)*. We treated two strains of *P. aeruginosa*, one with no genetic alterations (expressing *pqsE(WT)*) and one harboring the *pqsE(S285W)* mutation at its native locus on the chromosome with multiple concentrations of either HJ1 or HJ5. For comparison, pyocyanin production was measured in a Δ*pqsE* strain, which produced virtually no pyocyanin. At up to 100 µM HJ1, no appreciable effect on pyocyanin production was observed in either the *pqsE(WT)* or *pqsE(S285W)* strain (**Figure 5a**). On the other hand, at 50 µM, HJ5 inhibited pyocyanin production by approximately 30% compared to the 0 µM (DMSO) treated WT strain, and at 100 µM there was a ∼37% decrease (**Figure 5b**). While this was an exciting finding, a similar decrease in pyocyanin production was observed when the *pqsE(S285W)* strain was treated with HJ5, suggesting that the effect of HJ5 on pyocyanin production is through a mechanism that is independent of PqsE and, likely, the QS network.

**Figure 5:**
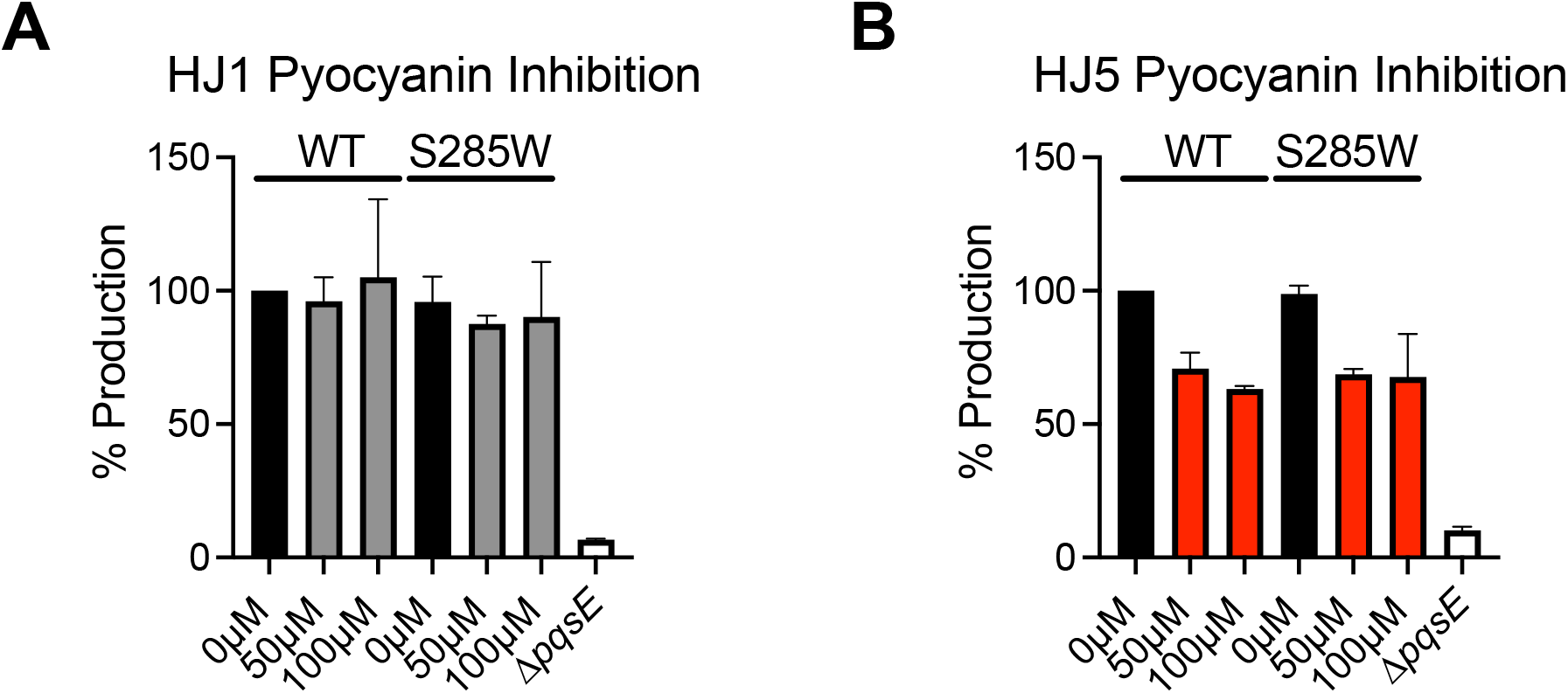
Neither HJ1 nor HJ5 inhibits pyocyanin production in a PqsE-specific manner. A) Effect of HJ1 on pyocyanin production in wildtype *P. aeruginosa* PA14 and a *pqsE(S285W)* mutant strain. Black bars represent pyocyanin production by the DMSO-treated cultures and grey bars represent cultures treated with HJ1 at either 50 or 100 µM. B) Effect of HJ5 on pyocyanin production in wildtype *P. aeruginosa* PA14 and a *pqsE(S285W)* mutant strain. Black bars represent pyocyanin production by the DMSO-treated cultures and red bars represent cultures treated with HJ5 at either 50 or 100 µM. Pyocyanin in the culture supernatants was measured as absorbance at 695 nm (OD_695_) and normalized to OD_600_ of the resuspended cell pellet. % Production is relative to the OD_695_/OD_600_ measured for DMSO-treated wildtype *P. aeruginosa*. Data shown are the average of biological duplicates and error bars represent standard deviation.

### HJ5 does not induce a conformational change in PqsE

To further investigate whether HJ5 is capable of inducing a conformational change in PqsE, we performed partial proteolysis using trypsin digestion. This technique has been used previously to observe differential digestion patterns that result from a protein being locked in alternate conformational states (28). We suspected that the PqsE(E182W) variant, harboring the described loop rearrangement, may yield a different banding pattern from PqsE(WT) upon a limited tryptic digest. Indeed, we found that following a 10-minute exposure to trypsin, the PqsE(E182W) variant displayed a unique band at ∼20 kDa that was not observed in the digested PqsE(WT) protein (**Figure 6a**, labelled “band 2”). We hypothesized that if a small molecule, such as HJ5, were able to bind in the active site of PqsE and induce a conformational change similar to the one characteristic of the PqsE(E182W) variant, we would expect to see the appearance of this ∼20 kDa band following treatment of the PqsE(WT)-inhibitor complex with trypsin. PqsE(WT) was treated with increasing concentrations of HJ5 prior to digestion with trypsin, and the resulting digests were analyzed by SDS-PAGE. Conclusively, treatment of PqsE(WT) with up to 100 µM HJ5 did not produce a banding pattern similar to that of PqsE(E182W) (**Figure 6a**). This suggests that HJ5 binding does not induce a conformational change, explaining the lack of effect on the PqsE-RhlR interaction and therefore virulence in *P. aeruginosa*. Notably, we observed that the band intensity of the undigested protein incubated with HJ5 increases as concentration of HJ5 increases (**Figure 6**, “band 1”). This result is consistent with the observation that HJ5 does bind to PqsE, and therefore appears to stabilize the protein upon exposure to trypsin. Analysis of the band intensities revealed that band 1 constitutes the majority of signal in the PqsE(WT) samples, whereas the digested PqsE(E182W) variant showed a decreased proportion of band 1 intensity and a corresponding increase in band 2 intensity (**Figure 6b**). These shifts further support the conclusion that while HJ5 binds to PqsE it does not mimic the effects of the E182W substitution on overall protein conformation.

**Figure 6:**
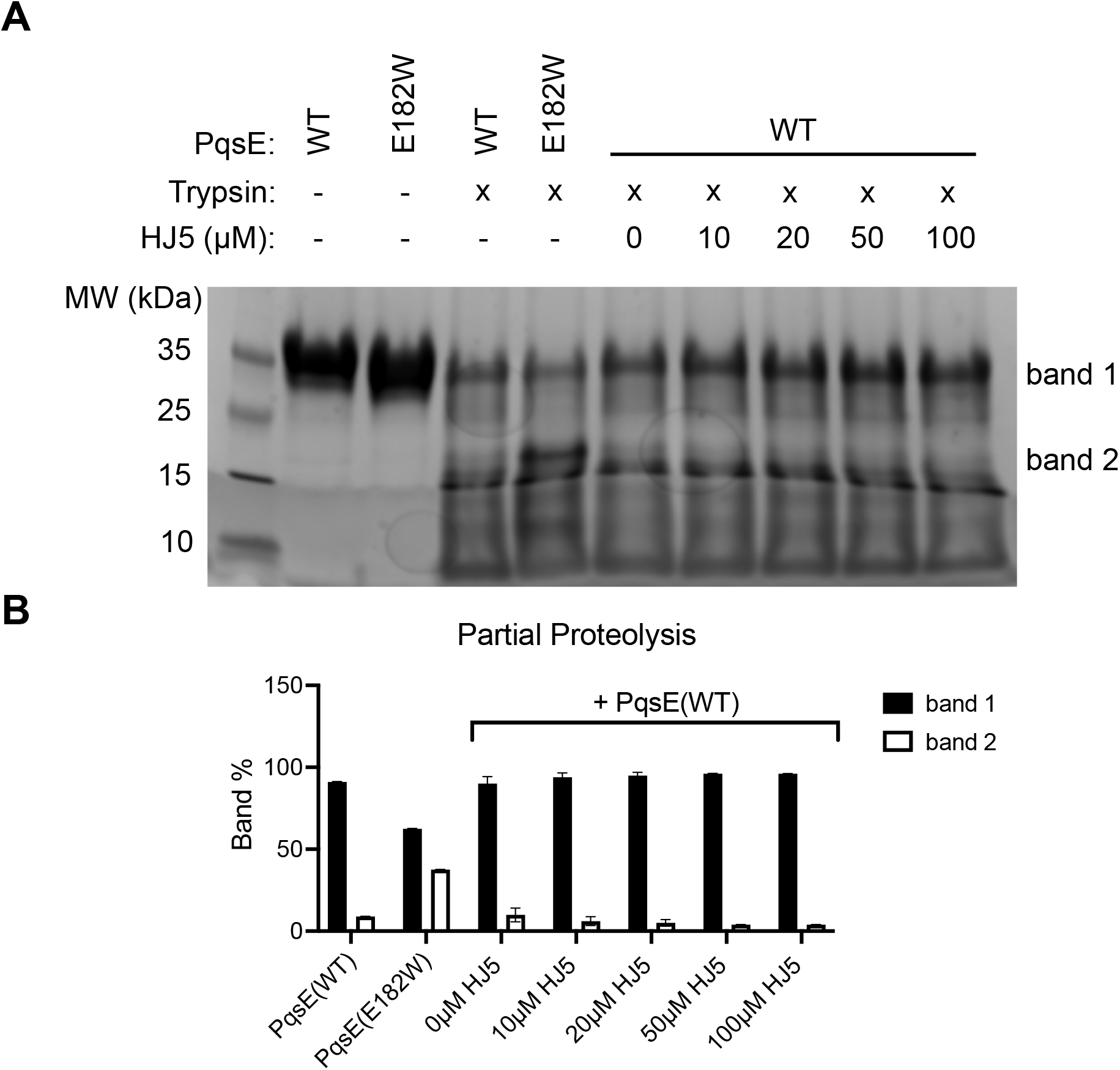
HJ5 does not induce a conformational change in PqsE. A) SDS-PAGE image of PqsE proteins partially digested with trypsin in the presence of varying concentrations of HJ5. In the first two lanes after the ladder, PqsE(WT) and PqsE(E182W) were not exposed to partial proteolysis by trypsin. The undigested protein runs as a single ∼34 kDa band (band 1). The next two lanes show the banding patterns of PqsE(WT) and PqsE(E182W) produced from a 10-min incubation with trypsin. The presence of a distinct band at ∼20 kDa (band 2) is diagnostic of the conformational shift induced by the E182W substitution. In the next 5 lanes, PqsE(WT) was incubated with the indicated concentrations of HJ5 prior to digestion by trypsin for 10 min. B) Quantification of bands produced by partial tryptic digest of PqsE. Only bands 1 and 2 were quantified and plotted as the % total band weight quantified per lane. The data shown in the graph represent all lanes of the gel in A, except for the first two lanes containing undigested protein. Values plotted are the average of duplicate experiments and error bars represent standard deviation. The second gel image used in this calculation is included in Supplemental Figure S3.

## Discussion

The PqsE-RhlR protein-protein interaction is currently under investigation as a promising target for new antimicrobial agents to treat *P. aeruginosa* infections (25, 29). Due to the complexity of targeting protein-protein interactions that occur over large, shallow protein surfaces (30, 31), it is particularly encouraging that mutagenic alterations in the active site of PqsE can allosterically affect the affinity for RhlR. Our hope is that a molecule can be designed to bind in the PqsE active site and induce the loop rearrangement characteristic of the PqsE(E182W) variant. While the molecules presented in this study do not achieve this effect, we have identified HJ5 as an active site-binder that has an altered mode of binding to PqsE. This is suggested by the incomplete inhibition profile of HJ5 compared to its parent scaffold, HJ1. We also gather that HJ5 likely has a differential mode of binding to PqsE from the shift in T_m_ induced by HJ5 upon binding to the PqsE(E182W) variant compared to HJ1 (**Figure S2**). This shift was not statistically significant compared to that observed for the HJ2 derivative, but it was greater than the shift observed for HJ1, which was a reversal of the trends observed for every other variant included in the experiment and was a consistent effect upon replicate experiments. Unfortunately, the PqsE(E182W) variant is severely impaired itself for both enzymatic capacity and driving pyocyanin production (through the weakened interaction with RhlR). Therefore, follow-on experiments measuring the effect of HJ5 binding PqsE(E182W) are limited. We are currently exploring other methods for studying this binding event.

In addition to presenting new PqsE inhibitors, we also present new methods for assessing activity of these molecules. Excitingly, the HJ series were completely incapable of binding the PqsE(S285W) variant. This was largely due to the loss of hydrogen bonding with the substitution of the serine, as binding affinity was also dramatically reduced for the PqsE(S285A) variant as well. This complete loss of binding allowed us to use a strain of *P. aeruginosa* expressing *pqsE(S285W)* as a tool strain to determine whether any *in vivo* effects of HJ1 or HJ5 on virulence were due to their effect on the PqsE-RhlR system. We have made earlier attempts to use this strain as a test of mechanistic specificity, however, previous attempts have been complicated by the finding that other molecules still bound PqsE(S285W), but with reduced affinity. The complete loss of inhibition of PqsE(S285W) by HJ1 and HJ5 made the results of *in vivo* specificity testing unambiguous.

Another new method introduced in this study for assessing PqsE conformational states is the use of partial proteolytic digest by trypsin. We found that the PqsE(E182W) variant, which is locked in an altered conformation with reduced affinity for RhlR, has a different banding pattern from PqsE(WT) upon limited digestion with trypsin. We used this partial proteolysis assay to show that, although HJ5 does bind PqsE (the increase in intensity of band 1 in the resulting gel supported this finding) it did not induce a conformational change, as there was no appearance of the unique PqsE(E182W) band (band 2). Our goal is to identify a molecule that induces a conformational change in PqsE similar to that observed in the PqsE(E182W) variant, and therefore, this type of analysis will be valuable for continued medicinal chemistry and drug discovery efforts.

## Experimental Procedures

### Strains and Media

All *P. aeruginosa* experiments were done using the UCBPP-PA14 strain background (referred to as PA14). All strains used in this study are listed in the Supporting Information (**Table S1**). All cultures were grown in Luria-Bertani (LB) broth (Difco). PqsE was recombinantly expressed and purified as described previously (18, 21).

### General Synthesis

Compounds HJ1-5 were all prepared via amide coupling reactions catalyzed by *N*-(3-Dimethylaminopropyl)-*N’*-ethylcarbodiimide (EDC) and 1-hydroxybenzotriazole (HOBt). The starting carboxylic acids and amines were purchased from VWR and Sigma Aldrich, with the exception of the starting carboxylic acid used to synthesize HJ5, which had to be prepared by the following method. All products were characterized by ^1^H NMR on a Bruker 400 MHz NMR as well as IR to confirm the loss of an amine peak, and appearance of an amide peak.

### Preparation of the carboxylic acid for HJ5

To prepare the carboxylic acid for HJ5 synthesis, 1 molar equivalent of methyl 2-amino-3,4-dimethylbenzoate and 2.3 molar equivalents of acetic anhydride were dissolved in chloroform and stirred at room temperature for 1 hour. 2.2 molar equivalents of isopentyl nitrite and 0.3 eq of potassium acetate were added and the reaction was heated to reflux for 24 hours. The reaction was then cooled, washed with sodium bicarbonate, and dried over sodium sulfate. The resulting ester was deprotected by adding 6 mL methanol and 4 mL 6 M HCl. The mixture was allowed to stir at room temperature for 18 h, after which the organic solvent was removed by rotary evaporation leaving an oil. The oil was triturated with ethyl acetate, vacuum filtered, and then washed twice with ethyl acetate. The resulting carboxylic acid was used without further purification.

### Synthesis of HJ1-HJ5

1 molar equivalent of a carboxylic acid-derivatized indazole was added to 1 molar equivalent of *t*-butyl (4-aminophenyl)carbamate and dissolved in DMF. Then 1.2 molar equivalents of both EDC and HOBt were added with stirring at room temperature for 8 hours until completion. The reaction was monitored by thin layer chromatography (TLC). Once complete, the reaction was washed with water and dried by vacuum filtration. The solid product was washed three times with ethyl acetate and then once with acetonitrile. The isolated product was determined to be pure by ^1^H NMR and was used without further purification.

### Compound Characterization

#### HJ1

^1^H NMR (400 MHz, DMSO) 13.09 (s, 1H), 10.31 (s, 1H), 9.25 (s, 1H), 8.16 (s, 1H), 8.01-7.99 (t, J = 5.6 Hz, 2H), 7.65-7.62 (d, J = 8.8 Hz, 2H), 7.40-7.38 (d, J = 8.4 Hz, 2H), 7.25-7.22 (t, J = 8.0 Hz, 1H), 1.43 (s, 9H).

#### HJ2

^1^H NMR (400 MHz, DMSO) 12.73 (s, 1H), 10.25 (s, 1H), 9.31 (s, 1H), 8.03-8.01 (d, J = 6.8 Hz, 1H), 7.97-7.92 (t, J = 12 Hz, 1H), 7.69-7.67 (d, J = 9.2 Hz, 2 H), 7.44-7.42 (d, J = 8.4 Hz, 2H), 7.22-7.11 (m, 1H), 3.32 (s, 3H) 1.31 (s, 9H).

#### HJ5

^1^H NMR (400 MHz, DMSO) 13.12 (s, 1H), 10.20 (s, 1H), 9.30 (s, 1H) 8.20 (s, 1H), 8.02-7.96 (m, 1H), 7.69-7.65 (t, J = 8.8 Hz, 3H), 7.46-7.41 (t, J = 9.8 Hz, 3H), 7.06-7.01 (t, J = 12.4 Hz, 1H), 3.31 (s, 3H), 1.47 (s, 9H).

### Esterase Assay

Purified PqsE, diluted in assay buffer (50 mM Tricine, 0.01% Triton X-100, pH 8.5), was added to the wells of an opaque 384-well plate at a final concentration of 125 nM. Test molecules were diluted in DMSO and added to the wells at the indicated concentrations (5% DMSO final). The plate was then incubated at room temperature for 5 minutes. MU-butyrate in assay buffer was added to the plate at a final concentration of 2 µM. The plate was incubated at room temperature for 20 minutes prior to measuring fluorescence (Excitation: 360 nm, Emission: 450 nm) to monitor the hydrolysis of MU-butyrate. All data were plotted in the Prism 9.0 program and IC_50_ values were calculated from inhibition curves fit to the data.

### Differential Scanning Fluorimetry

Compounds HJ1, HJ2, and HJ5 were tested for the ability to bind and shift the melting temperature (T_m_) of purified PqsE and PqsE variants using differential scanning fluorimetry (DSF) as described previously with some modifications (18). Briefly, compounds in DMSO were diluted in assay buffer (50 mM Tris, 150 mM NaCl, 200 µM MnCl_2_, pH 8) to achieve a final concentration of 100 µM (1% DMSO). Compound solutions were transferred to the wells of a 96-well PCR plate (ThermoFisher) and control wells were prepared with 1% DMSO (negative control). SYPRO Orange Dye and PqsE solutions were diluted in assay buffer and subsequently added to the wells to reach final concentrations of 7x SYPRO Orange and 2 µM PqsE. Final volume in each well was 80 µL. Plates were sealed with transparent optical adhesive covers (ThermoFisher) prior to measurement of SYPRO Orange fluorescence (ex: 580 nm, em: 623 nm) in a QuantStudio 3 Real-Time PCR system (ThermoFisher) over an increasing temperature gradient from 25 to 90 ºC (temperature increased at a rate of 0.1 °C/s). T_m_ was determined from first derivative plots of the raw fluorescence curves using the Prism 9.0 software.

### Pyocyanin Assay

*P. aeruginosa* PA14 cultures were grown overnight in LB at 37 °C with shaking. The saturated cultures were then diluted 1:1000 into fresh LB containing test molecules in DMSO (final 1% DMSO). Cultures then grew for ∼17 hours at 37 °C with shaking at 200 rpm. Next, 1 mL of each culture was collected and pelleted by centrifugation at 13,000 rpm for 3 minutes. The supernatant was carefully transferred to a cuvette, and the OD_695_ was measured in a Genesys 20 ThermoSpectronic spectrophotometer. The cell pellet was resuspended in 1 mL of PBS (137 mM NaCl, 2.7 mM KCl, 10 mM Na_2_HPO_4_ 1.8 mM KH_2_PO_4_, pH 7.4) and then transferred into the wells of a clear 96-well plate at 100 µL per well. The OD_600_ was measured to assess cell density. Pyocyanin production (OD_695_) of the supernatants was normalized to cell density (OD_600_ of the resuspended cell pellets).

### Partial Proteolysis

All proteins were thawed on ice prior to use. PqsE(WT) or PqsE(E182W) was aliquoted into microcentrifuge tubes and diluted to a final concentration of 10 µM in partial proteolysis buffer (50 mM Tris, 150 mM NaCl, pH 8). To each reaction, 1 µL of DMSO or HJ5 at varying concentrations is added. Reactions were incubated for 10 minutes at room temperature. Proteolysis was initiated by the addition of 10 µL of 0.25% trypsin, and proteolytic digest was allowed to occur for 10 minutes at room temperature. Reactions were then quenched with the addition of 33 µL 4x Lamelli dye and boiled at 95 ºC for 3 minutes. The samples were then subjected to SDS-PAGE and the resulting gel images were analyzed in the Bio-Rad Image Lab software.

## Data Availability

Data generated in this study are contained within the article and are available from the corresponding author (I.R.T.) upon request.

## Supporting Information

Additional inhibition curves, DSF data, a replicate gel, and a table of all strains used in this study are included in the supporting information (PDF).

## Acknowledgments

We thank members of the Taylor group for helpful advice and discussions.

## Author Contributions

S.B.O., H.A.J., M.G.O., and K.R.S. conducted experiments and curated data. S.B.O., H.A.J., M.G.O., K.R.S., and I.R.T. designed experiments and prepared the manuscript.

## Conflict of Interest

The authors have no conflicts of interest to declare.

## Notes

### Competing Interest Statement

The authors have declared no competing interest.

