## Supplemental figures and tables for "New inhibitors of the *Pseudomonas aeruginosa* enzyme, PqsE, and methods assessing their potential to induce a conformational change via active site binding"

**This PDF file includes:**

Figure S1
Figure S2
Figure S3
Table S1

**Supplementary Figures**

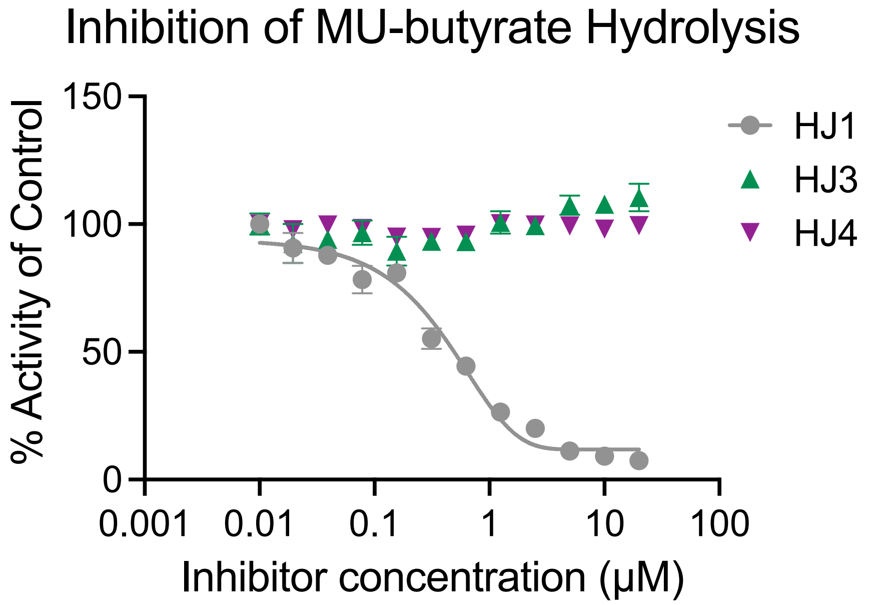


**Figure S1:** HJ2 and HJ3 do not inhibit PqsE enzymatic activity. Inhibition of PqsE-catalyzed MU-butyrate hydrolysis is shown. For comparison, HJ1 inhibits PqsE esterase activity with an $IC_{50}$ of ~400 nM, whereas the chlorinated and brominated derivatives, HJ3 (green triangles) and HJ4 (purple upside-down triangles) show no inhibitory activity. Background fluorescence was measured for the compound dilution series plus MU-butyrate in the absence of PqsE and was subtracted from the values plotted prior to normalization. The data were normalized so that DMSO-treated PqsE is 100 % activity. Data shown are the average of technical triplicates and error bars represent standard deviation.

**A**

### $T_m$ of PqsE-HJ complexes

| | DMSO | HJ1 | HJ2 | HJ5 |
|---|---|---|---|---|
| WT | 66.9 | 70.5 | 67.7 | 68.8 |
| E280A | 66.5 | 68.8 | 67.1 | 67.7 |
| E182W | 73.4 | 74.4 | 73.9 | 75.5 |
| E182W/E280A | 69.8 | 72.4 | 70.0 | 71.9 |
| S285W | 66.7 | 66.8 | 66.8 | 67.5 |
| S285A | 67.3 | 68.1 | 67.9 | 68.6 |
| E182A | 69.6 | 70.1 | 69.2 | 70.6 |

**B**

### $\Delta T_m$ of PqsE-HJ complexes

| | DMSO | HJ1 | HJ2 | HJ5 |
|---|---|---|---|---|
| WT | 0.00 | 3.56 | 0.74 | 1.82 |
| E280A | 0.00 | 2.28 | 0.61 | 1.21 |
| E182W | 0.00 | 0.98 | 0.48 | 2.05 |
| E182W/E280A | 0.00 | 2.58 | 0.20 | 2.12 |
| S285W | 0.00 | 0.07 | 0.07 | 0.74 |
| S285A | 0.00 | 0.81 | 0.61 | 1.35 |
| E182A | 0.00 | 0.50 | -0.47 | 1.00 |

**Fig S2**: $T_m$ measurements for PqsE(WT) and PqsE variants in the presence of HJ derivatives. PqsE (2 μM) was incubated with each molecule (100 μM) and the resulting $T_m$ was measured by DSF. A) All average $T_m$ values determined from three independent experiments performed in triplicate. Color scale is set with the lowest value in the whole data set (66.5) as white and the highest value (75.5) as green. B) The $\Delta T_m$ was determined by subtracting the DMSO-treated protein $T_m$ from each inhibitor-treated protein $T_m$. As in A, the color scale is set with the lowest value in the whole data set (-0.47) as white and the highest value (3.56) as green.

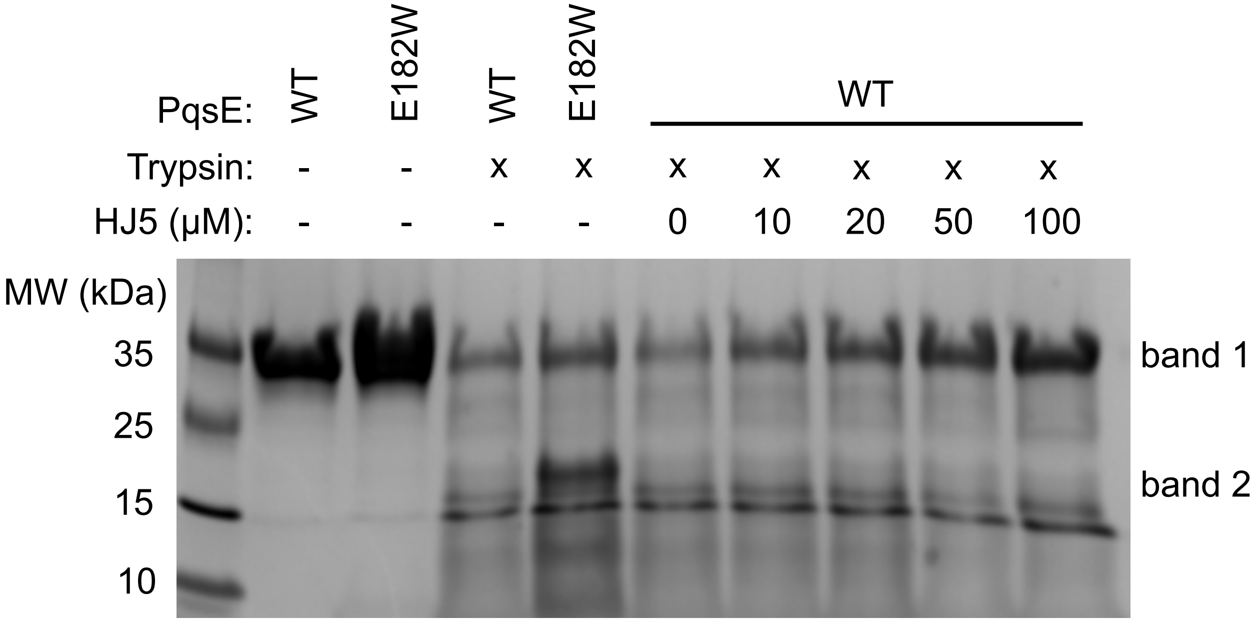


**Fig S3:** Additional partial proteolysis experiment. SDS-PAGE image of PqsE proteins partially digested with trypsin in the presence of varying concentrations of HJ5. In the first two lanes after the ladder, PqsE(WT) and PqsE(E182W) were not exposed to partial proteolysis by trypsin. The undigested protein runs as a single ~34 kDa band (band 1). The next two lanes show the banding patterns of PqsE(WT) and PqsE(E182W) produced from a 10-min incubation with trypsin. The presence of a distinct band at ~20 kDa (band 2) is diagnostic of the conformational shift induced by the E182W substitution. In the next 5 lanes, PqsE(WT) was incubated with the indicated concentrations of HJ5 prior to digestion by trypsin for 10 min. Band weight calculations were performed on this gel image and used along with the gel image in Fig 6a to generate the data shown in Fig 6b.

**Supplementary Tables**

| **Strain** | **Description** | **Reference** |
|---|---|---|
| UCBPP-PA14 | PA14 *P. aeruginosa* Wildtype | Laboratory stock |
| SM776 | *E. coli* BL21 (DE3) pET28b-6xHis-pqsE(WT) | 22 |
| IT39 | *E. coli* BL21 (DE3) pET28b-6xHis-pqsE(S285A) | 18 |
| IT55 | *E. coli* BL21 (DE3) pET28b-6xHis-pqsE(S285W) | 18 |
| IT105 | PA14 *P. aeruginosa* Δ*pqsE* | 21 |
| IT169 | PA14 *P. aeruginosa pqsE(S285W)* | This study |

**Table S1.** Strains used in this study.
